# Extraordinary claims require extraordinary evidence in the case of asserted mtDNA biparental inheritance

**DOI:** 10.1101/585752

**Authors:** Antonio Salas, Sebastian Schönherr, Hans-Jürgen Bandelt, Alberto Gómez-Carballa, Hansi Weissensteiner

## Abstract

A breakthrough article published in PNAS by Luo et al. (2018) challenges a central dogma in biology which states that the mitochondrial DNA (mtDNA) is inherited exclusively from the mother. By sequencing the mitogenomes of several members of three independent families, the authors inferred an unprecedented pattern of biparental inheritance that requires the participation of an autosomal nuclear factor in the molecular process. However, a comprehensive analysis of their data reveals a number of issues that must be carefully addressed before challenging the current paradigm. Unfortunately, the methods section lacks any description of sample management, validation of their results in independent laboratories was deficient, and the reported findings have been observed at a frequency at complete variance with established evidence. Moreover, the remarkably high (and unusually homogeneous) levels of heteroplasmy reported can be readily detected using classical techniques for DNA sequencing. By reassessing the raw sequencing data with an alternative computational pipeline, we report strong correlation to the NextGENe results provided by the authors on a per sample base. However, the sequencing replicates from the same donors show aberrations in the variants detected that need further investigation to exclude contributions from other sources or methodological artifacts. Finally, applying the principle of *reductio ad absurdum*, we demonstrate that the nuclear factor invoked by the authors would need to be extraordinarily complex and precise in order to preclude linear accumulation of mtDNA lineages across generations. We discuss alternate scenarios that explain findings of the same nature as reported by Luo et al., in the context of *in-vitro* fertilization and therapeutic mtDNA replacement ooplasmic transplantation.

## Claims on biparental inheritance

Luo et al. (2018) describe apparent biparental inheritance patterns of mitochondrial genomes (mtDNA) in three different families. A total of 21 samples were reported, some in replicates, and sequenced by different laboratories using various methods to verify the results and rule out the possibility of contamination. Based on the findings, the authors describe an unprecedented pattern of autosomal dominant-like inheritance mode of the paternal mtDNA and propose the existence of a nuclear factor necessary in the molecular process. In some family members, the inherited “paternal” mtDNA exceeds the amount of maternal mtDNA. In addition, paternal mtDNA levels are excessively high in all reported cases.

However, there are a number of serious gaps regarding description of the methodology employed and the interpretation of the findings. By any measure, the results are extraordinary, sharply contrasting with the extensive and well-established experience of the scientific community specializing in mtDNA research.

Confirmation of substantial germline contributions of paternal mtDNA would be of significant consequence, not only because it contradicts a well-established tenet of biology, namely, strict maternal inheritance of the human mtDNA genome, but also because many disciplines of science are affected, including, molecular anthropology, population genetics, and clinical and forensic genetics. The article by Luo et al. (2018) has received international attention in the media, as well as critical and supporting commentaries (summarized in **Supplementary Info**).

## Methodological issues in Luo’s study

According to Kallet (2004), a methods section *“requires a precise description of how an experiment was done…The description of preparations, measurements, and the protocol should be organized chronologically.*” The methods are at least as important as the results themselves, and this is particularly true when findings have the impact of those claimed by Luo et al. (2018). However, significant issues in the article by Luo et al. (2018) need clarification. First, information on sample collection, handling and preparation was completely omitted, in addition to specifications on where and how the analyses (sequencing, Restriction Fragment Length Polymorphism typing [RFLP], etc.) were carried out at the different laboratories involved (**Figure S1; Supplementary Info**). Second, information on sequencing replicates is limited and contradictory. Third, sequencing replicates show unexpected differences in the reported variants and/or heteroplasmic levels (**Figure S2**, **S3**, **S4**, **and S5;** see **Supplementary Info**). These differences are so significant that they could even be attributed to different donors (according to e.g. guidelines of the International Society of Forensic Genetics; ISFG (Parson *et al.*, 2014)). Fourth, the sequencing results summarized in three key figures have been deficiently and ambiguously reported. Detailed concerns regarding the methodology are provided in **Supplementary Info**.

### Improbably frequent incidences of the phenomenon

Luo et al. (2018) found initial evidence of biparental mtDNA inheritance after testing their first proband (IV-2 of Family A), and other family members (all carrying high levels of heteroplasmy). Next, they examined two other families, solely applying the selection criteria of suspected mitochondrial involvement (but never confirmed in the present analysis): “*to confirm the findings in Family A, we recruited two additional families (Families B and C)…*” (Luo et al., 2018). It seems that the authors did not need extensive sampling efforts to detect new candidate families, because, their next two searches simply yielded the same extraordinary findings observed in Family A (that is, three out of three exhibited the same exceedingly rare trait of biparental mtDNA inheritance). Two of the families were recruited in the MitoClinic and another one in their collaborative Mayo Clinic laboratory which indicates that their findings are a relatively common phenomenon (e.g. roughly as frequent as many low frequency “Mendelian” diseases). The high prevalence of the phenomenon in the sampling population with no evidence of mtDNA pathology is inconsistent with the authors’ use of the term “unusual”, which only refers to the fact that the phenomenon has not been reported before (with a single exception, discussed below).

The described findings are extraordinary insofar as they are in sharp conflict with current evidence: there are thousands of laboratories working on mtDNA in different fields of research; and hundreds of thousands of haplotypes reported, not only in databases (e.g. so far over 47,000 whole mtDNA sequences and 71,000 control region sequences have been deposited in GenBank by January 1st, 2019), and in the literature (probably triplicating the numbers in databases), but also obtained by widely used private companies e.g. offering direct-to-consumer-tests. Given this large amount of mtDNA sequence data, the very high levels of heteroplasmy reported could not have remained undetected even with the classical procedures of DNA sequencing (e.g. Sanger-based). The senior author in Luo et al. (2018), Dr. Taosheng Huang, has in fact recognized the ability of Sanger to detect heteroplasmies above a threshold of 20% (or even smaller) (Tang & Huang, 2010). Therefore, with heteroplasmy levels in the mixtures with a claimed paternal contribution of 24% to 76%, classical Sanger-based sequences would suffice to detect these levels. The additional resolution of NGS or of third-generation sequence is not necessary to detect such high levels of heteroplasmy (see for instance, Figure 3B of Luo et al. (2018)). The fact that the claimed phenomenon of biparental inheritance has not been observed before by other researchers for more than three decades is odd, particularly in areas of research with special emphasis on methodological quality and high technical standards typified by forensic genetics (**Supplementary Info**).

Taking into account the above issues, the argument of the authors that “*clinical laboratories that perform such analyses tend to report only previously reported pathogenic or likely pathogenic mutations. Unusual results are often ignored, especially when they do not involve likely pathogenic variants*” seems indefensible (a shared interpretation with others (McWilliams & Suomalainen, 2019)). Such arguments question the wide-scale professional activity carried out in thousands of clinical laboratories worldwide, as well as in thousands of institutions working in molecular anthropology and forensic genetics.

The frequency of the findings described by Luo et al. (2018) cannot be compared with the unique pathological case described in the literature on human paternal inheritance (Schwartz & Vissing, 2002). This earlier case of biparental mtDNA inheritance (see the doubts raised by Bandelt et al. (2005)), was noted by Luo et al. (2018), but unfortunately, their comments suffer from literature bias (**Supplementary Info**).

### A sophisticated ‘molecular surgery’

Luo et al. (2018) propose the existence of a nuclear factor that would explain the observed pattern of biparental mtDNA inheritance. To understand how this nuclear factor could explain the observed pattern of inheritance, let us first examine the pedigree of Family A more deeply.

The authors initially deduce (no data are available) the mtDNA genome haplotypes of the first generation (I-1 and I-10) based on the haplotype mixture found in the next generations. With this assumption in mind, the daughter II-4 would inherit a mixture of haplogroups: R0a1 (∼39%) from father (I-1) and H1a1 (∼60%) from mother (I-10) (**Figure 1A**). Then, individual III-6 -daughter of II-4-would inherit from II-4 the R0a1 haplotype exclusively; surprisingly, at almost the same proportion (∼40%), while receiving 60% of U5b1d1c from the mother (II-40). In other words, their paternal (II-4) H1a1 mtDNAs disappeared completely and are replaced exactly by their maternal (II-40) mtDNAs (**Figure 1A**). Such highly precise “molecular surgery” is not mentioned in the text at all: the authors just speculate about the possible existence of a nuclear DNA (nDNA) factor that is inherited in a dominant way; and such a dominant nDNA factor would necessarily be transmitted from males I-1 > to II-4; only to end in female III-6. However, to fully explain the inheritance pattern reported, this nDNA factor should give response to both (*i*) the described biparental inheritance of the mtDNA, and (*ii*) the complete elimination in the progeny of all mtDNAs that do not have a strict ancestral paternal origin in all previous generations. Thus, it is necessary to invoke an extremely complex molecular mechanism that specifically targets the patrilineal or the matrilineal inherited mtDNA of these biparental mtDNA males such that the only haplotypes that are transmitted (again, at unusually high frequency) to the next generation are those strictly inherited from carrier males. Exactly the same phenomenon occurs in Family B (**Figure 1B**), and Family C (**Figure 1C**).

**Figure 1.**
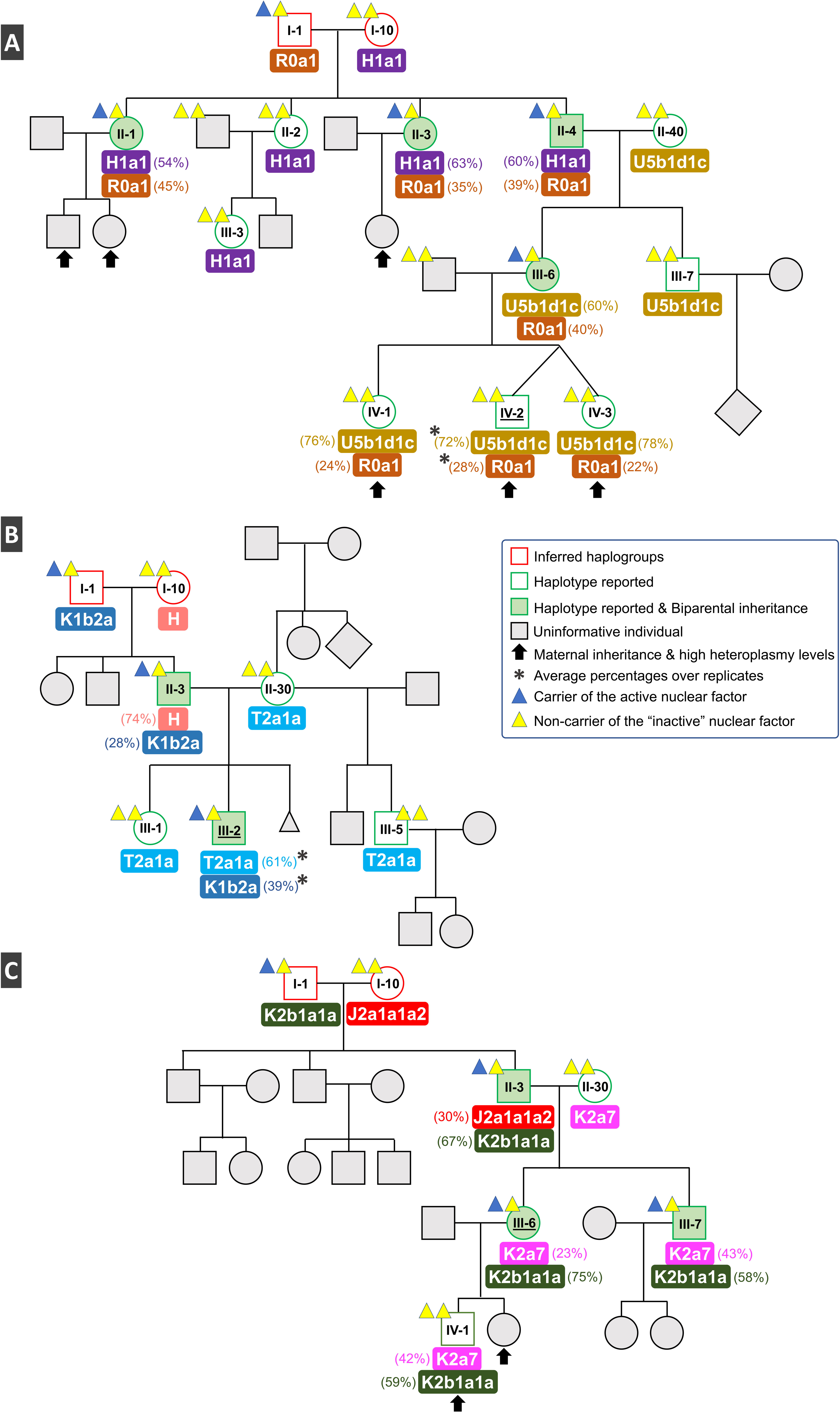
Reconstruction of pedigrees (Figure 1A, 1B, and 1C relate to Family A, B and C, respectively) reported in Figure 1A, 2A and 3A from Luo et al. (Luo et al., 2018) indicating the most remarkable issues from this publication. The percentages of heteroplasmies indicated correspond to the averages computed from their supplementary files: percentages of heteroplasmies are taken from the diagnostic positions for each haplogroup and averaged; these percentages never sum to 100% in the reported data. We added to the pedigree the dominant nuclear factor that, according to the authors, would necessary be invoked to explain the patterns of biparental inheritance. Note also that heteroplasmic fathers (e.g. II-4 Family A; H1a1+R0a1), only transmit to the siblings (e.g. III-6 Family A) mtDNAs originally obtained through the patriline (e.g. I-1 Family A; R0a1), so the ones inherited from the matriline (I-10 Family A; H1a1) disappear in the descendants (III-6 Family A). The “molecular surgery” needed to explain this pattern should be highly precise; this observation has been omitted by the authors. The colored triangles indicate the most parsimonious pattern of transmission of the nuclear factor that would better fit the inheritance patterns observed, a fact that is ambiguously described by the authors. The grey polygons indicate parts of the pedigrees shown by the authors but that are uninformative. The index cases were underlined.

An interesting consequence of this “molecular surgery” mechanism would be to preclude the progressive lineal accumulation of an undefined number of haplotypes through generations; e.g. without this mechanism, III-6 should carry a combination of three haplotypes (H1a1+R0a1+U5b1d1c) instead of the two reported (R0a1+U5b1d1c).

In addition, given that this nuclear factor resides in the autosomes, women would be potential carriers of such a genetic factor (if inherited from a paternal carrier; e.g. III-6). However, the authors do not offer any explanation whether such factors can be somehow activated or inactivated for future generations (e.g. in their male descendants). The most plausible explanation is that the nuclear factor does not play a role when carried by the mothers; if so, some kind of e.g. epigenetic control should be invoked to explain the fact that only male carriers have the ability to produce biparental inheritance.

The most intriguing consideration comes from the possibility that a non-carrier (of the nuclear factor) heteroplasmic woman (e.g. IV-1, and IV-3 from family A; with heteroplasmy pattern U5b1d1c + R0a1) will at some point, have descendants with male carriers of the nuclear factor (e.g. III-2 from family B; **Figure S6A**). The same would occur if we assume that the heteroplasmic woman is a carrier (**Figure S6B**). Then, we cannot abandon the possibility to find individuals with an undetermined number of different mtDNA haplotypes. Unless another extraordinary molecular ability is added to the nuclear factor; yet such *ad hoc* patterns remain to be reported in the literature and databases. It can be additionally argued that there have been plenty of opportunities during human evolutionary history to give rise to such multi-haplotypic (multi-heteroplasmic) male and female individuals.

Finally, another remarkable feature of the data is not only the unusual homogeneous heteroplasmic levels observed in the pedigrees, but also that these levels are not necessarily explained by the presence of the nDNA factor invoked by the authors. For instance, individuals IV-1, IV-2, and IV-3 all inherited virtually the same heteroplasmic combination (∼75% U5b1d1c + ∼25% of R0a1) from their mother (III-6).

### About alternate scenarios of apparent biparental inheritance

There are some conceivable scenarios that could generate signals of apparent biparental inheritance; that is, the existence of more than one haplotype coexisting in the same individual with at least two different donors:

- The simplest explanation is the accidental mixing of blood samples (or DNA, PCR amplicons, etc), extracted from different donors on the bench. The senior author in Luo et al. (2018) has already tested the ability of massively parallel sequencing using artificial mixtures of samples combined at different molar ratios; see Tang and Huang (2010). Extensive literature exists describing errors due to unintended mixtures during the mtDNA analysis process (**Supplementary Info**). Co-authors in the Luo et al. (2018) article from Baylor laboratory (Zhang *et al.*, 2012) have experience of this problem using almost the same methodology as that described in Luo et al. (2018).
- Single long-range PCR. Paradoxically the long-range PCR covering the complete mitochondrial genome, intended to avoid co-amplification of nuclear encoded mitochondrial pseudogenes (NuMTs) could be too unspecific, due to primer problems leading to co-amplification of NuMTs (Luo *et al.*, 2019, Lutz-Bonengel & Parson, 2019). Therefore, the mtDNA to nDNA ratio could be observed at the presented high levels.
- Bone marrow transplantation or other types of transplant might give rise to chimerisms that mimic a pattern of biparental inheritance. The forensic field has already reported scenarios where simple Sanger sequencing has detected such patterns (Seo *et al.*, 2012).
- Mitochondrial genome editing *in vivo* made to correct pathogenic mtDNA variation (Gammage *et al.*, 2018) could also mimic scenarios of seeming biparental inheritance.
- Several techniques of *in vitro* fertilization (IVF) using therapeutic mtDNA replacement (MRT) have been developed and have experienced substantial advances in the last few years (Rai *et al.*, 2018). As early as 1999, Brenner et al. (2000) described the presence of heteroplasmy after human ooplasmic transplantation (a technique developed in the late 90’s). In 2000, St John et al. (2000) described the detection of paternal mtDNA after IVF and intracytoplasmic sperm injection (ICSI) treatment. In 2001, Barritt et al. (2001) stated that ooplasmic transfer from fertile donor oocytes into patient oocytes led to the birth of about 30 children. These authors reported that heteroplasmy in the blood from healthy children originated from ooplasmic transfers. More recently, Kang et al. (2016), with the participation of coauthors in Luo et al. (2018), have described the use of spindle transference of mtDNA in human oocytes, which involved an efficient replacement (>99%) of oocyte donor’s mtDNA. Authors in Luo et al. (2018) have also played a leading role in the pioneering study reporting the first live three-parent baby birth derived from oocyte spindle transfer with the aim of preventing mitochondrial disease (Slone *et al.*, 2017, Zhang *et al.*, 2017). In addition, the senior author in Luo et al. (2018), has recently proposed scenarios of “*Forced inheritance of paternal mtDNA: a potential alternative for MRT*” as a plausible alternative to MRT to prevent the transmission of mtDNA-related diseases (Slone et al., 2017). It is worth mentioning that haplogroup matching should be considered for mitochondrial transplantation according to the latest HFEA (Human Fertilisation and Embryology Authority) panel guidelines. The panel further recommends that “*haplotype information on the recipient and the donor is recorded*”. However due to the diversity of mitochondrial haplotypes, haplogroup matching is not always successful in preventing low-level heteroplasmies or mito-nuclear incompatibilities (Morrow *et al.*, 2015).

None of these potential scenarios were described by Luo et al. (2018) in their tested families. However, all laboratories investigating this phenomenon should be aware of these possible artificial signals of biparental mtDNA inheritance (especially cases emerging from the increasing number of three-parent babies as a result of MRT) as well as other sources of biparental mtDNA inheritance or similar confounding situations such as artificial amplification of nuclear encoded mitochondrial pseudogenes (NuMTs) or artifacts generated by long range PCR. In this regard, it is most paradoxical that a few individuals have large deletions at significant frequencies (e.g. individual A-II-2 has a deletion from positions 2010 to 2205 in >46% of the mtDNAs; **Figure S7** and **Supplementary Info**). This observation has a possible technical origin (e.g. artifacts related to long range PCR) and could partly explain the haplotype differences observed between replicates (e.g. those affecting annealing primer regions).

## Conclusions

That mtDNA is transmitted from the mother to the offspring constitutes a central tenet of biology for decades. Therefore, claims of biparental inheritance of the mtDNA require exceptional evidence (Deming, 2016, Truzzi, 1978), and such evidence should come from fully independent laboratories, not just those involved in the original publication. Validation should be based on the use of original blood samples from donors; analyses based on amplicons are insufficient and cannot validate the biological process *per se* but validate only a final technical step of the whole process. Aiming to facilitate such independent validation, we were unsuccessful in obtaining key biological samples from the authors of Luo et al. (2018), because they denied our request for blood samples.

Therefore, serious open questions remain in the reported work of Luo et al. (2018) that need to be clarified before accepting these results as evidence of biparental inheritance (*contra* (Vissing, 2019)). Likewise, it is important for journals to demand full and complete evidence from authors that challenge well-established paradigms. Further work is needed before proper assessment can be made of the extraordinary claim of paternal mtDNA inheritance.

## Acknowledgements

We thank Chris Phillips and Bernhard Rupp for critically reading this paper, and Lukas Forer for useful and important discussions.

## Author contribution

All the authors contributed to the analysis of the data and writing of the manuscript.

## Supplementary file

**Figure S1.** Indication of the laboratories where the different sequencing analysis were carried out according to the information extracted from the data provided by the authors upon request.

**Figure S2.** Haplotype differences in sequencing replicates as read from FASTQ files sent by the authors upon request and including the three replicates reported by the authors, namely, “(blood)”, “(blood) rpt1”, and “(blood) rpt2” (here re-labeled as R1, R2, and R3, respectively). Position 310 was eliminated from comparisons as it was not reported by the authors.

**Figure S3.** Variation in heteroplasmic levels reassessed with mtDNA-Server (https://mtdna-server.uibk.ac.at; (Weissensteiner *et al.*, 2016)), in the index patients of Family A (A-IV-2) and Family B (B-III-2) where replicates where carried out. The stacked barplots represent the mean heteroplasmic levels of the haplotype mixture, with the different colors denoting the corresponding haplogroup. Sample A-IV-2 is a mixture of U5b1d1c and R0a1 with the latter being considered the paternal mtDNA contribution, while B-III-2 is a mixture of T2a1a with K1b2a with the latter considered as the paternal contribution.

**Figure S4.** Comparison of variant calling pipeline in sample A-IV-2. The raw data obtained from the authors was analyzed with mtDNA-Server yielding to highly concordant results to the NextGENe data also provided by the authors. The plot denotes the major haplotype (heteroplasmy level ≥50%) and minor haplotype (heteroplasmy level <50%). The replicate A-IV-2_CCHMC_3 shows a switched major haplotype compared to the other replicates.

**Figure S5.** Comparison of variant calling pipeline in Sample B-III-2. The raw data obtained from the authors was analyzed with mtDNA-Server not available for all replicates, however yielding to highly concordant results to the NextGENe data. Two replicates (B-III-2_Baylor and B-III-2_HUANG_3) present noise in the NextGENe data, attributable to homopolymeric stretch around position 309 on the rCRS in B-III-2_Baylor and heteroplasmic mutations around 2,700 in B-III_HUANG_3). A significant fluctuation between the replicates can be observed.

**Figure S6.** We reproduce a hypothetical pedigree where a woman with the same heteroplasmic characteristics as IV-1 or IV-3 have descendants with a male carrying the nuclear factor, e.g. as III-2 from Family B (who is also heteroplasmic). The possibility of the woman being a carrier of the (activated/inactivated?) nuclear factor (B) or not (A) cannot be disregarded. To simplify the scenario, assume that this nuclear factor is transmitted by women in inactivated status. Several genetic conformations would be possible for the descendants; note that according to the patterns shown by the authors, it would be possible to observe descendants with three (or any number in other hypothetical scenarios) distinct haplogroups (e.g. d-2 in [A] and d-2 and d-3 in [B]). Note that, according to the extraordinary characteristics of the molecular surgery mechanisms proposed in the text, it is necessary to assume that only the mtDNA strictly inherited from the father of III-2 (haplogroup K1b2a) is transmitted to the progeny.

**Figure S7.** Analysis of deletions on eKLIPse of mtDNA FASTQ files provided by the authors upon request.

