## Supplementary figures and images for "Extraordinary claims require extraordinary evidence in the case of asserted mtDNA biparental inheritance"

### Figure S1

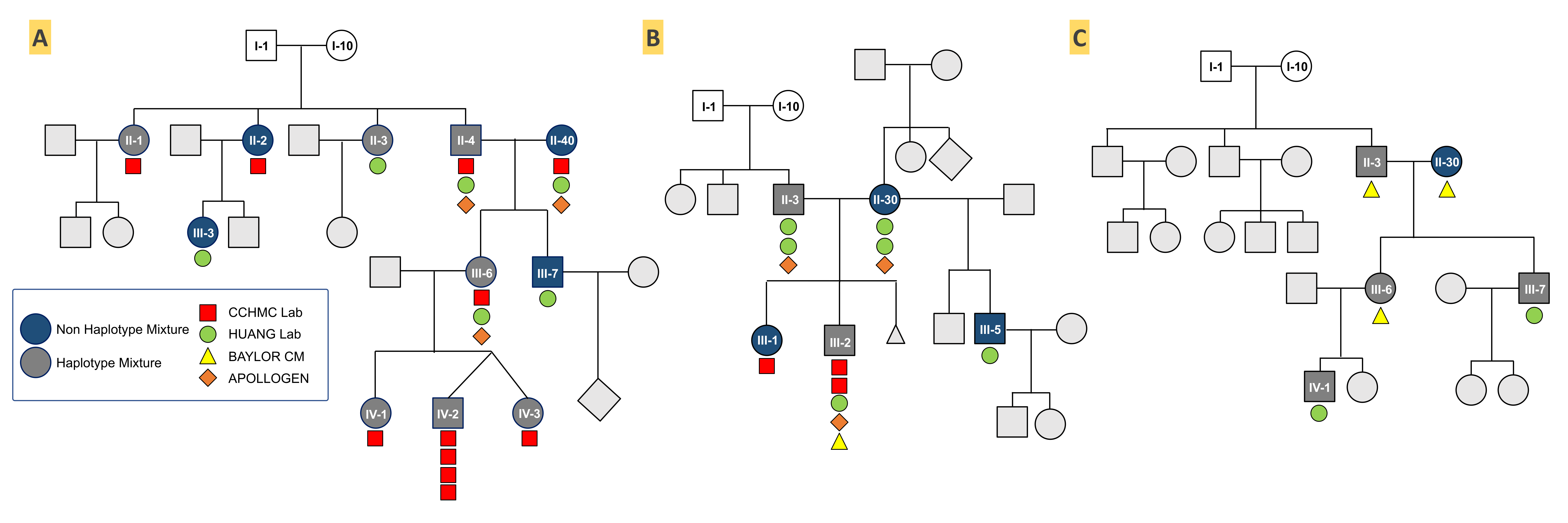

### Figure S2

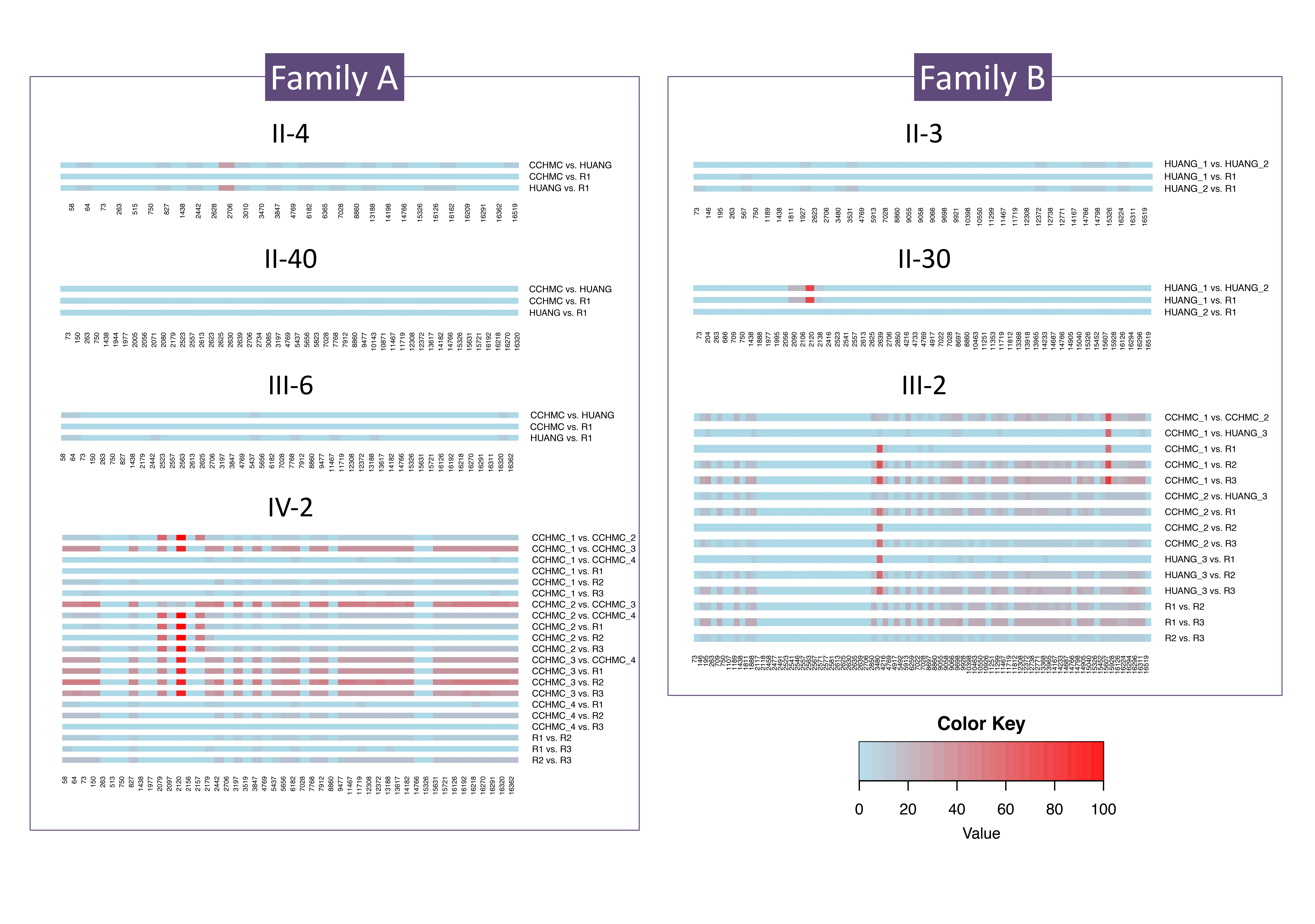

### Figure S3

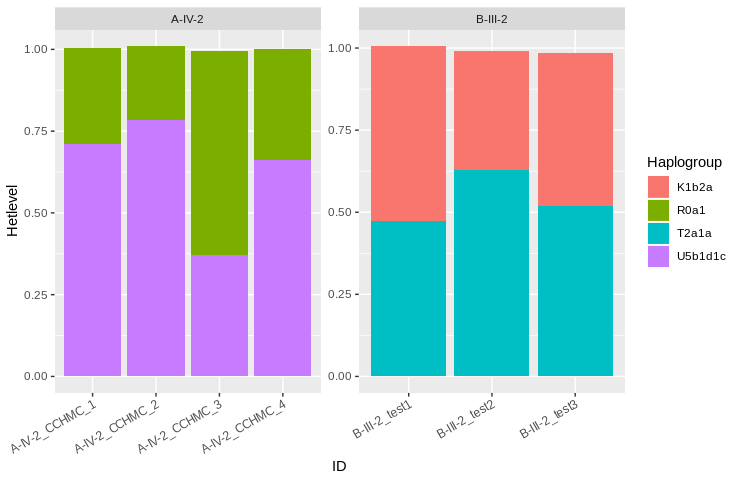

### Figure S4

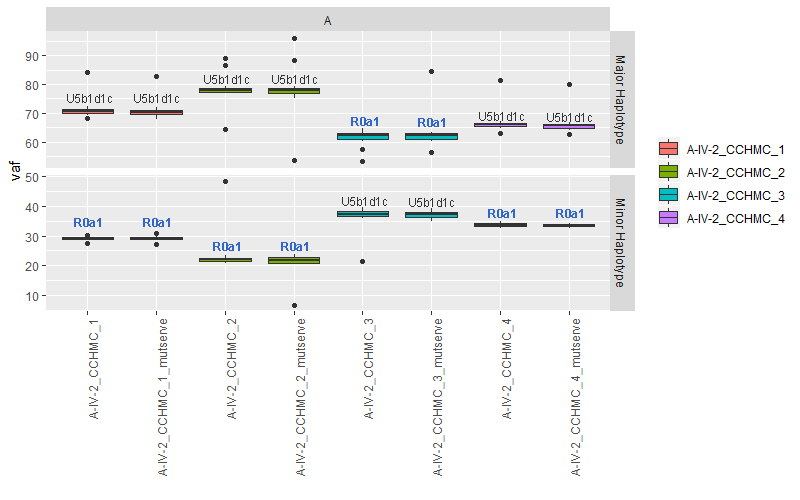

### Figure S5

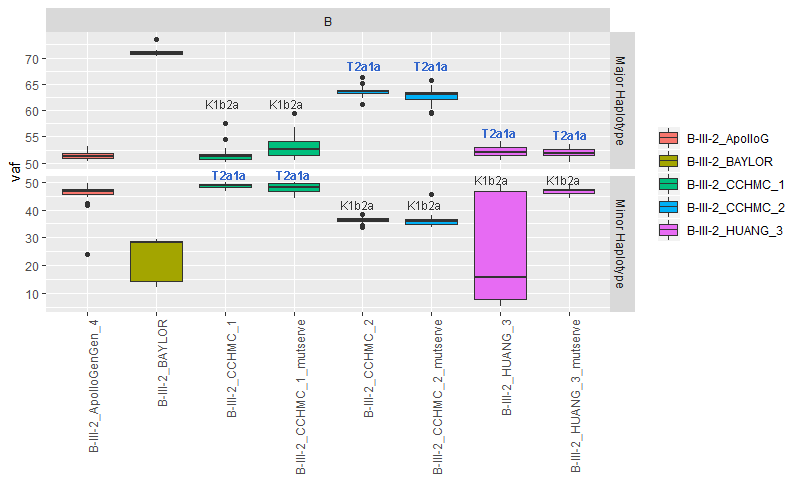

### Figure S6

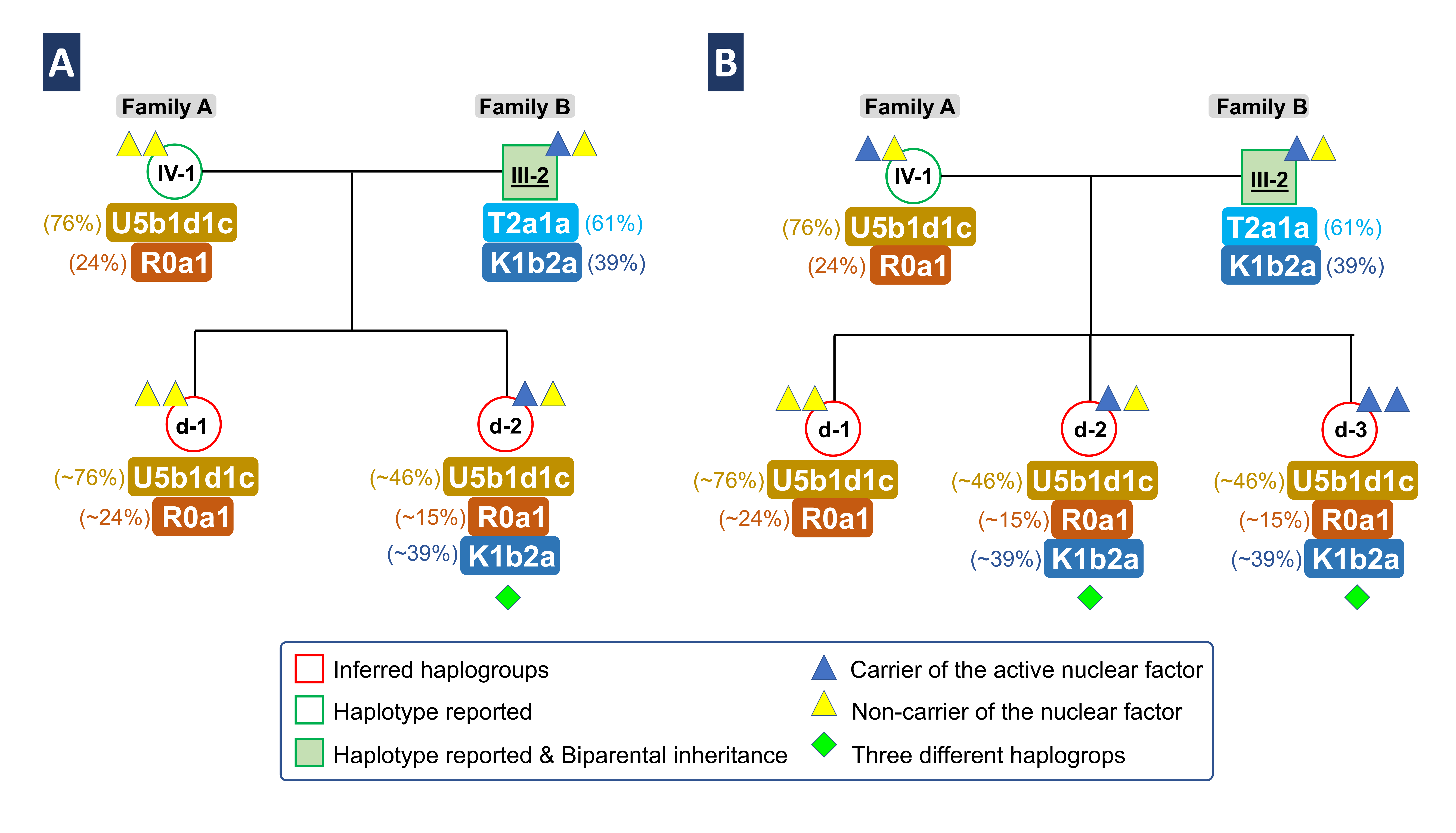

### Figure S7

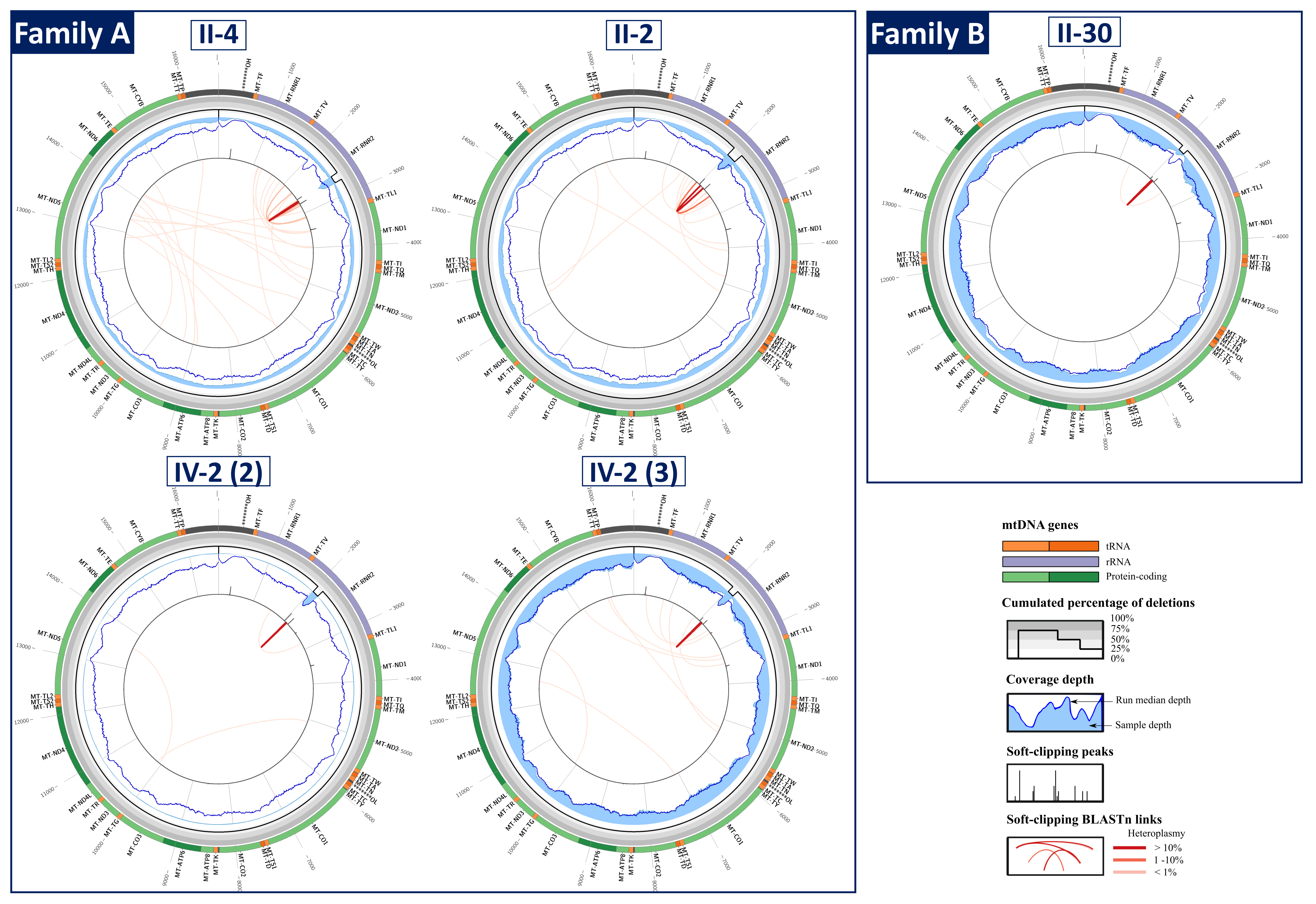
