## Supplementary Text for "Extraordinary claims require extraordinary evidence in the case of asserted mtDNA biparental inheritance"

**1. Recent comments / letters on the topic**

Recently two reviews/comments in *Proceedings of the National Academy of Sciences* (PNAS)(Vissing, 2019) and one in Nature (McWilliams & Suomalainen, 2019), address the paper of Luo et al. (2018) and its possible impact on mtDNA research.

In his review in PNAS, Vissing - co-author of the 2002 report in NEJM together with Schwartz, where both describe the unique case of biparental mtDNA inheritance (Schwartz & Vissing, 2002) - addresses the impact of mtDNA inheritance on genetic counseling, as well as forensic science and anthropology. He classifies the results as convincing, since it was reported in three families by using “whole-DNA” sequencing. To be correct – it should be noted that only whole mtDNA sequencing was performed, ignoring the nuclear portion of the genome. However, he noticed a “puzzling finding in the paper by Luo et al”, pointing to the uniform level of heteroplasmy in individuals in the same family. As an explanation he indicates the possibility of co-amplification of nuclear encoded mitochondrial pseudogenes (NuMTs).

This is also one of the central points by Lutz-Bonengel’s and Parson’s letter in PNAS (Lutz-Bonengel & Parson, 2019), indicating the lack of methods to exclude the possibilities to sequence NuMTs. Furthermore, the lack of father’s data, which was deduced based on the mixture model, cannot assert the paternal inheritance. In their reply, Luo et al. (2019) state that they did not propose where the mtDNA sequences were from. According to the authors, the presence of NuMTs is extremely unlikely as they could observe 100 copies of mtDNA genomes per nuclear genome.

McWilliams and Suomalainen (2019) report on the “provocative study” by Luo et al. (2018), in a comment in *Nature*. Their review notes the differences in the mitochondrial content between human eggs (100,000×) and sperm (100×), without addressing the topic of NuMTs. However, the question of why this phenomenon of the biparental mtDNA inheritance remained undetected, arises. McWilliams and Suomalainen (2019) see an unsatisfactory explanation by the authors, who argue that it was often overlooked, as no disease causing variant is involved in the mixture of haplotypes. Furthermore, the review highlights the lack of causal link to disease but underlines the importance of the findings for mitochondrial replacement therapy.

The representation in McWilliams and Suomalainen’s Figure 1, that visualizes our previous concerns about the mixture of three different haplotypes that should be expected, while only two are detected, remains uncommented.

**2. Methodological issues in Luo’s et al.**

There are methodological issues that need clarification by the authors:

2.1. Information omitted from Materials and Methods

The publication contains only two sections on Materials and Methods: one briefly describing the patients, and another describing sequencing methods. The information on sample collection, handling and preparation is missing. Issues on where and when some procedures were carried out on sampling management add important uncertainty on the procedures used. For instance, it is unclear if the fresh blood samples were always managed by the same laboratory/researcher.

2.2. Confused handling of sequencing replicates

It is unclear where and how the replicates were carried out based on the publication alone, although one can infer from their text and their supplementary information data that only the index cases of family A (IV-2) and B (III-2) were replicated (IV-2 apparently by the same lab [CCHMC] and III-2 in another independent laboratory not indicated). Family C was analyzed in the Diagnostic Laboratory at Baylor College of Medicine (no replications were mentioned in the article). Moreover, there is no information on where each of the three replicates was processed and sequenced (in the case of sample III-2 from Family B). It is not clear if the sequencing results reported correspond to the first round of sequencing, or those generated by SMRT in their ‘verification’ of the findings.

More details on the laboratories involved in the sequencing analyses have been obtained from the authors upon request, and this information is summarized in **Figure S1**. This information adds much more potential for variation to the overall scenario because:

(a) the authors mention two clinics where the samples were recruited and (seemingly) handled (MitoClinic, and Mayo Clinic), and three laboratories responsible for the sequencing analyses, namely, CCHMC and the Baylor laboratory and the Genome Science University of Maryland laboratory. However, in the data sent by the authors there are four laboratories mentioned: CCHMC, Baylor, Apollogen, and Huang.

(b) data from PacBio was omitted;

(c) the responsible laboratories for the different analyses do not fit with the information described in the text, e.g. in the methods section it is said that Family A and B were analyzed in CCHMC, while Family C in Baylor (see contradiction in **Figure S1**);

(d) the authors only provided partial raw data, so it is not possible to evaluate the reproducibility of the findings in all laboratories.

(e) Sample chronologies obtained from FASTQ file creation dates do not fit with the chronology described by the authors; e.g. they indicated that Family A was the first to be recruited and analyzed, however FASTQ files of individuals III-1 and III-2 from Family B dated from 2014, that is, two years before the generation of the FASTQ files from Family A.

**Figure S1** also indicates that the experimental design for sequencing replications and validation is inconsistent; e.g. only sample III-2 was replicated in four laboratories, while e.g. individuals from Family C were sequenced only once.

2.3. Information on the biological source used for independent verification of sequencing results is missing

Verification of the sequencing results was carried out in different laboratories with a different sequencing technology (SMRT with PacBio) and using long range PCR products of the entire mtDNA. Information on the biological source used for each verification is missing. Using the same PCR amplicons for the samples would allow the possibility of error carryover from previous steps. Overall, sequencing results were not adequately validated, and samples cannot be considered to be fully independent replicates.

2.4. Haplotype differences in replicates

Parallel sequencing carried out on samples extracted from the same donor should yield exactly the same haplotypes, or at most, very minor differences in heteroplasmic levels (arising from limitations of the sequencing chemistries and sequencers). Strikingly, sequencing replicates in Luo et al. do not show consistent results; against Luo et al. (2019) e.g. “*…these results were fully consistent with the previous sequencing*” and Vissing (2019) e.g. “*…the findings are convincing because it was demonstrated in three independent families, in several generations of these families, and by using whole-DNA sequencing performed by two different laboratories*”).

We have reanalyzed the FASTQ files provided by the authors upon our request. For this purpose, we generated FastQC reports, mapped the files with BWA MEM 0.7.17-r1188, generated QualiMap 2 (Okonechnikov *et al.*, 2016) reports by using MultiQC (Ewels *et al.*, 2016) for a summarize analysis of both FastQC and QualiMap 2 data, and performed the variant calling with a local version of mtDNA-Server (Weissensteiner *et al.*, 2016) (v.1.1.17 see https://github.com/genepi/mutserve), with the default parameters (including BAQ, per base quality 20, alignment quality 30 and mapping quality 20) (Li, 2011). We further compared the NextGENe data provided by the authors as Text files and Excel files to our results within R. We noticed issues around the primer sites (i.e. 2120), but also around 2625, 3480 or 10283) in the NextGENe data, which otherwise strongly correlated with our results. **Figure S2** represents the haplotypes differences from the replicates, where the variant level varies significantly between the runs. The barplots of **Figure S3** show variation in heteroplasmic levels reassessed with mtDNA-Server in the index patients of Family A and Family B. Given the high accuracy of NGS, the contributions from parents fluctuate remarkable in both samples depending on the laboratories and replicates. When further exploring the heteroplasmic levels of sample A-IV-2 using other variant calling pipelines, we observe an unexpected switch of the major haplotype compared to the other replicates (**Figure S4**). A similar pattern is observed for the index case B-III-2 (**Figure S5**) together with a significant fluctuation between the replicates.

The results of cross-comparisons of all replicates contrasted with the high sensitivity, specificity and accuracy of parallel sequencing reported only a few years ago by Tang and Huang (2010) (Huang, senior author in Luo et al. (2019)); an article reporting experiments on different ratios of human mtDNA mixtures. Overall, sequencing replicates have so many differences (**Figure S2**) that they could be attributed to different, unrelated donors. This is difficult to understand given the high sequence coverage observed in the corresponding FASTQ files (above 9,000× in all the samples).

2.5. RFLP data missing

Additional RFLP experiments on selected individuals were carried out; but no information on the laboratory involved nor on which source the PCR amplification was performed, is provided in the publication.

2.6. SNP data

As described by the authors, 14 nuclear markers of the same sample were “genotyped” using “either TaqMan assay or Sanger sequencing” (which is not a genotyping technique per se) in order to trace possible cross-contamination during procedures. Information on the markers used (e.g. their rs identification numbers) was not provided.

2.7. Data from some of the individuals analyzed are missing

The authors reported high levels of heteroplasmy in three individuals of Family A (III-1, III-2; III-5) and 1 individual of Family C (IV-2) but the results were not reported, but only inferred from the maternal heteroplasmy present.

2.8. Defective figures

The three key figures in the publication contain ambiguous information. To give an example from Family A, e.g. the group of samples II-1, II-3, and II-4 or the group III-6, IV-1, IV-2, IV-3 have similar heteroplasmy levels. However, the heteroplasmic proportions displayed in the histograms from Figure 1B, and the values reported in their table from Figure 1C, are not explained and how they were rounded to 100%. It is necessary to imagine that some kind of average over variants was computed; but these averages do not necessarily fit with the values reported in the supplementary files (which are never rounded to 100%).

For heteroplasmic individuals, there are homoplasmic variants for which their parental origin cannot be differentiated from the data. For instance, in Figure 1B, they reported the same (average?) profile for samples IV-1, IV-2, IV-3 and III-6; that is a mixture of R0a1 plus U5b1d1c (inherited from their heteroplasmic mother III-6). However, R0a1 and U5b1d1c share common variants (e.g. at positions 263, 750, etc) for which it is not possible to infer if they proceed from R0a1 or from U5b1d1c haplotypes. Therefore, it is important to clarify that the information on these variants, although not necessarily wrong, is also derived from the data.

2.9. Highly homogeneous intergenerational heteroplasmic levels

The authors realized that for Family A, the inherited mutations over three generations are “oddly coincidental”, without investigating any further. Family A index patient IV-2, and his siblings, are the result of, at most, grand-paternal mitochondrial contribution. However, the grandparent was a mixture of two haplotypes where exactly one haplotype got inherited to the mother of the index patient, mixed with the haplotype of the grandmother. It is further “odd” that the heteroplasmy levels remain the same for the deduced paternal haplotype R0a1, although mixed with two different haplotypes across the pedigree tree.

2.10. Batch processing

As denoted by the authors, whole mtDNA sequencing was carried out in batches of at least 12 samples in one pool, then loaded on to the sequencer. It is not clear if all sequencing facilities used the same protocol.

In brief, description of procedures has important gaps in the Material and Methods section, and it seems that validation was carried out only on selected individuals and on amplicons previously generated by CCHMC and the Diagnostic Laboratory at Baylor College of Medicine.

**Literature bias**

The paper by Luo et al. (2018) also suffers from bias in the literature selection. They cited Schwartz and Vising (2002) and its follow-up study (Kraytsberg *et al.*, 2004), where a unique case of paternal inheritance (plus mtDNA recombination) was reported. However, Luo et al. (2018) fail to mention that Bandelt et al. (2005b) already raised serious questions with respect to these earlier findings. Surprisingly, the recent article by Slone et al. (2017), with its three authors co-authoring Luo et al. (2018) and the same corresponding author, recognize that “*there is considerable debate about the accuracy and interpretation*” of the Schwartz and Vissing (2002) findings.

**Extraordinary claims**

Due to the special nature of the casework carried out in forensic genetic laboratories (a key service in the justice system of most countries worldwide), it is a priority to pay particular attention to potential errors in mtDNA casework and accompanying reference databases. To give an example, the reference mtDNA forensic international database EMPOP (https://empop.online; (Parson & Dür, 2007)) has implemented a number of procedures, described in the literature, that allow the detection of errors before the sequences are entered in the database. When potential sequencing errors or anomalies are detected by EMPOP, these are further explored by the laboratories involved and, after further examination of the original samples, the errors are confirmed and repaired. Any unusual finding (e.g. evidence for biparental inheritance) would be made public due to the importance of such phenomenon to forensic casework. Exhaustive validation exercises related to the mtDNA test are also common activities in international forensic institutions, e.g. (Alonso *et al.*, 2002, Crespillo *et al.*, 2006, Montesino *et al.*, 2007, Prieto *et al.*, 2003, Salas *et al.*, 2005b). Unless we assume an exceptional nature of biparental inheritance (in full contrast to the findings of Luo et al. (2018)), it is not credible that this phenomenon has passed undetected by forensic experts.

In addition, there are several dozen articles in the literature where a large number of errors in mtDNA databases have been reported e.g. (Achilli *et al.*, 2008, Bandelt *et al.*, 2005a, Bandelt *et al.*, 2007, Bandelt *et al.*, 2002a, Bandelt & Salas, 2009, Bandelt & Salas, 2012, Bandelt *et al.*, 2004a, Bandelt *et al.*, 2006, Bandelt *et al.*, 2009, Bandelt *et al.*, 2002b, Bandelt *et al.*, 2004b, Kong *et al.*, 2008, Kong *et al.*, 2006, Salas *et al.*, 2007, Salas *et al.*, 2005a, Salas *et al.*, 2014, Salas *et al.*, 2005c, Yao *et al.*, 2008, Yao *et al.*, 2009) (a fact not mentioned by Luo et al. (2018)). The great majority of such errors still lack replies from the authors of the affected publications. In this regard, the criticism by Luo et al. (2018) of the study of Bandelt et al. (2005b), where many of these errors were revealed, are vague and unfounded. Last but not least, such comments could encourage other laboratories to relax necessary quality controls.

**Detection of large mitogenome deletions**

We ran eKLIPse (Goudenege *et al.*, 2018) for the detection of deletions in the FASTQ files obtained from the authors. It is interesting that a few individuals have large deletions affecting most of the mitogenome and at high frequency, namely, A-II-2 (~46%), A-II-4 (“Huang” replicate; ~34%), A-IV-2 (replicates 2 and 3 in CCHMC; ~34% and ~32% respectively), and B-II-30 (“Huang” replicate; ~26%); **Figure S7**.

It is noteworthy that these deletions have 5’ and 3’ breakpoints located at gene 16S rRNA (3’ breakpoints at positions 2010, 2026, 2028, and 2572; 5’ breakpoints at positions 2154, 2156, 2205, and 2715). It is probably not coincidental that these breakpoints correspond to the primer annealing regions reported by the authors in their Method 1, namely, Forward-2120 and Reverse-2019. This association cannot be established for their Method 2 because the corresponding FASTQ files were not provided by the Baylor laboratory.
